# Leveraging prior concept learning improves ability to generalize from few examples in computational models of human object recognition

**DOI:** 10.1101/2020.02.18.944702

**Authors:** Joshua S. Rule, Maximilian Riesenhuber

## Abstract

Humans quickly learn new visual concepts from sparse data, sometimes just a single example. Decades of prior work have established the hierarchical organization of the ventral visual stream as key to this ability. Computational work has shown that networks which hierarchically pool afferents across scales and positions can achieve human-like object recognition performance and predict human neural activity. Prior computational work has also reused previously acquired features to efficiently learn novel recognition tasks. These approaches, however, require magnitudes of order more examples than human learners and only reuse intermediate features at the object level or below. None has attempted to reuse extremely high-level visual features capturing entire visual concepts. We used a benchmark deep learning model of object recognition to show that leveraging prior learning at the concept level leads to vastly improved abilities to learn from few examples. These results suggest computational techniques for learning even more efficiently as well as neuroscientific experiments to better understand how the brain learns from sparse data. Most importantly, however, the model architecture provides a biologically plausible way to learn new visual concepts from a small number of examples, and makes several novel predictions regarding the neural bases of concept representations in the brain.

**Author summary:** We are motivated by the observation that people regularly learn new visual concepts from as little as one or two examples, far better than, e.g., current machine vision architectures. To understand the human visual system’s superior visual concept learning abilities, we used an approach inspired by computational models of object recognition which: 1) use deep neural networks to achieve human-like performance and predict human brain activity; and 2) reuse previous learning to efficiently master new visual concepts. These models, however, require many times more examples than human learners and, critically, reuse only low-level and intermediate information. None has attempted to reuse extremely high-level visual features (i.e., entire visual concepts). We used a neural network model of object recognition to show that reusing concept-level features leads to vastly improved abilities to learn from few examples. Our findings suggest techniques for future software models that could learn even more efficiently, as well as neuroscience experiments to better understand how people learn so quickly. Most importantly, however, our model provides a biologically plausible way to learn new visual concepts from a small number of examples.

## Introduction

Humans have the remarkable ability to quickly learn new concepts from sparse data. Preschoolers, for example, can acquire and use new words on the basis of sometimes just a single example [1], and adults can reliably discriminate and name new categories after just one or two training trials [2–4]. Given that principled generalization is impossible without leveraging prior knowledge [5], this impressive performance raises the question of how the brain might use prior knowledge to establish new concepts from such sparse data.

Several decades of anatomical, computational, and experimental work suggest that the brain builds a representation of the visual world by way of the ventral visual stream, along which information is processed by a simple-to-complex hierarchy, up to neurons in ventral temporal cortex that are selective for complex objects such as faces, objects and words [6]. According to computational models [7–11] as well as human functional magnetic resonance imaging (fMRI) and electroencephalography (EEG) studies [12,13], these object-selective neurons in high-level visual cortex can then provide input to downstream cortical areas, such as prefrontal cortex (PFC) and the anterior temporal lobe (ATL), to mediate the identification, discrimination, or categorization of stimuli, as well as more broadly throughout cortex for task-specific needs [14]. It is at this level where these theories of object categorization in the brain connect with influential theories of semantic cognition that have proposed that the ATL may act as a “semantic hub” [15], based on neuropsychological findings [16–18] and studies that have used fMRI [19–22] or intracranial EEG (iEEG) [23] to decode category representations in the anteroventral temporal lobe.

Computational work suggests that hierarchical structure is a key architectural feature of the ventral stream for flexibly learning novel recognition tasks [24]. For instance, the increasing tolerance to scaling and translation in progressively higher layers of the processing hierarchy due to pooling of afferents preferring the same feature across scales and positions supports robust learning of novel object recognition tasks by reducing the problem’s sample complexity [24]. Indeed, computational models based on this hierarchical structure, such as the HMAX model [25] and, more recently, convolutional neural network (CNN)-based approaches have been shown to achieve human-like performance in object recognition tasks [26–30], given sufficient numbers of training examples and even accurately predict human neural activity [31].

In addition to their invariance properties, the complex shape selectivity of intermediate features in the brain, e.g., in V4 or posterior inferotemporal cortex (IT), is thought to span a feature space well matched to the appearance of objects in the natural world [28,30]. Indeed, it has been shown that re-using the same intermediate features permits the efficient learning of novel recognition tasks [28,32–35], and the re-use of existing representations at different levels of the object processing hierarchy is at the core of models of hierarchical learning in the brain [36]. These theories and prior computational work are limited, however, to re-use of existing representations at the level of objects and below. Yet, as mentioned before, processing hierarchies in the brain do not end at the object-level but extend to the level of concepts and beyond, e.g., in the ATL, downstream from object-level representations in IT. These representations are importantly different from the earlier visual representations, generalizing between exemplars to support category-sensitive behavior at the expense of exemplar-specific details [37]. Intuitively, leveraging these previously learned visual *concept* representations could substantially facilitate the learning of novel concepts, along the lines of “a platypus looks a bit like a duck, a beaver, and a sea otter”. In fact, there is intriguing evidence that the brain might leverage existing concept representations to facilitate the learning of novel concepts: in *Fast Mapping* [1–3], a novel concept is inferred from a single example by contrasting it with a related but already known concept, both of which are relevant to answering some query. Fast Mapping is more generally consistent with the intuition that the relationships between concepts and categories are crucial to understanding the concepts themselves [38–41]. The brain’s ability to quickly master new visual categories, then, may depend on the size and scope of the bank of visual categories it has already mastered. Indeed, it has been posited that the brain’s ability to perform Fast Mapping might depend on its ability to relate the new knowledge to existing schemas in the ATL [42]. Yet, there is no computational demonstration that such leveraging of prior learning can indeed facilitate the learning of novel concepts. Showing that leveraging existing concept representations can dramatically reduce the number of examples needed to learn novel concepts would not only provide an explanation for the brain’s superior ability to learn novel concepts from few examples, but would also be of significant interest for artificial intelligence, given the current deep learning systems still require substantially more training examples to reach human-like performance [31,43].

We here show that leveraging prior learning at the concept level in a benchmark deep learning model of object recognition leads to vastly improved abilities to learn from few examples. We specifically find that broadly tuned conceptual representations can be used to learn visual concepts from as few as two positive examples, while visual representations from earlier in the visual hierarchy require significantly more examples to reach comparable levels of performance.

## Results

To explore whether concept-level leveraging of prior learning leads to superior ability to learn novel concepts compared to leveraging learning at lower levels, we conducted large-scale analyses using state-of-the-art CNNs (we also conducted similar analyses using the HMAX model [25,44], obtaining qualitatively similar results, albeit with overall lower performance levels). Specifically, we examined concept learning performance as a function of training examples for four feature sets (Conceptual, Generic_1_, Generic_2_, Generic_3_, Fig 1) extracted from a deep neural network (GoogLeNet [45]). Based on prior work using GoogLeNet, we hypothesize that the Conceptual features best model semantic cortex (e.g. ATL), while the Generic layers best model to high-level visual cortex (e.g. V4, IT, fusiform cortex) [30,31]. We predicted that higher levels would support improved generalization from few examples, and in particular that leveraging representations for previously learned concepts would strongly improve learning performance for few examples. To test this latter hypothesis, we modified the GoogLeNet architecture to perform 2,000-way classification. We then trained the modified network to recognize 2,000 concepts from ImageNet [46] (S1 Table). We examined the activations of each feature set for images drawn from 100 additional concepts from ImageNet (S2 Table), distinct from the previously learned 2,000 concepts.

**Fig 1.**
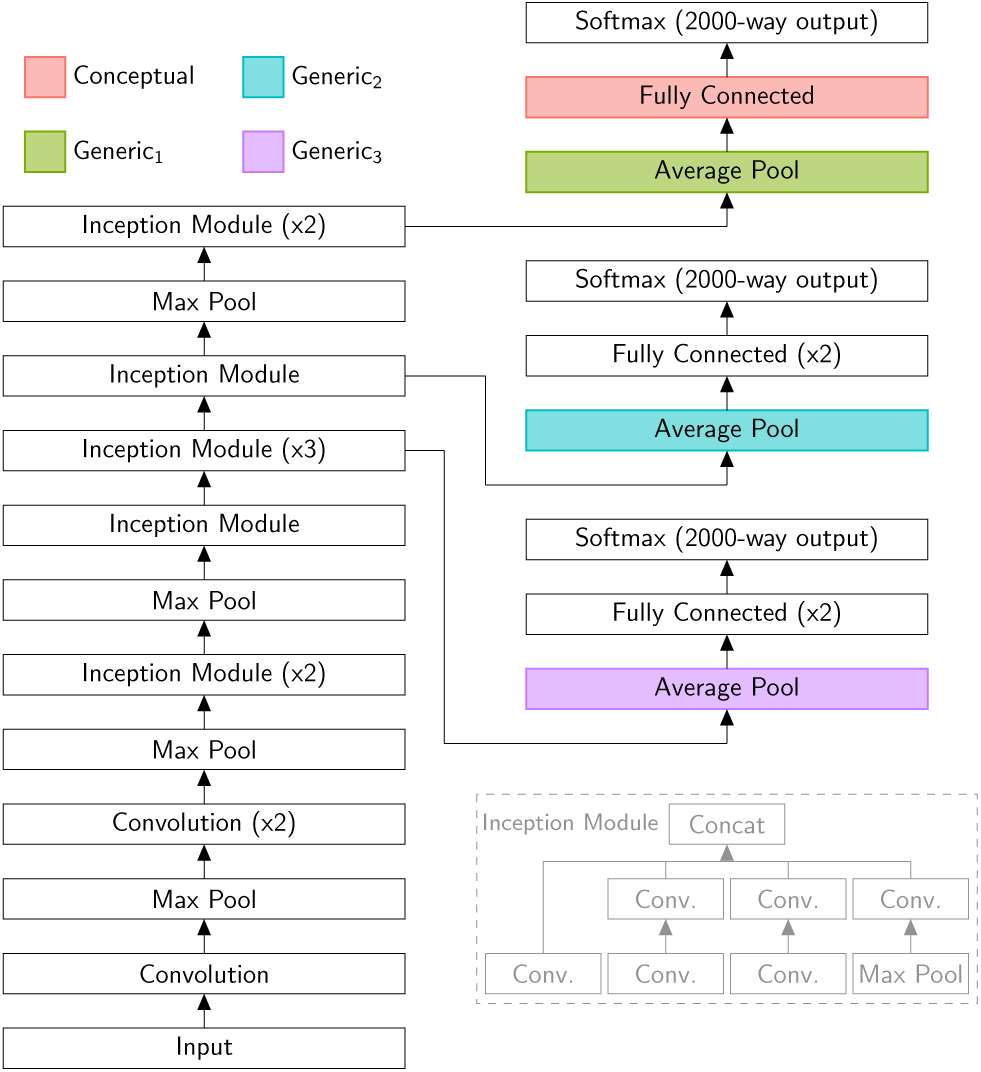
Network schematic. A schematic of the GoogLeNet neural network [45] as used in these simulations (main figure) and a schematic of the network’s Inception Module (gray inset on lower right). We modified the network to produce 2,000-way outputs, simulating representations for 2,000 previously learned categories. We then investigated how well representations at different levels of the hierarchy supported the learning of novel concepts. To encourage generalization, we wanted each layer to be broadly tuned, so we drew our conceptual layer not from the task-specific and sharply tuned final decision layer (Softmax), but the immediately preceding layer. Multiples (i.e. x2 or x3) indicate several identical layers being connected in series.

For our scheme to work, conceptual features must support generalization by being broadly tuned. All the feature sets we analyzed are thus part of the standard GoogLeNet architecture and come before the network’s final decision layer. The binary classifiers we trained for this analysis, however, were separate from GoogLeNet. We do not claim that they are part of the visual hierarchy so much as we use them to assess the usefulness of different parts of that hierarchy for sample-efficient learning.

The concepts GoogLeNet learns are based on visual information only and therefore do not capture the fullness of the rich and nuanced concepts used in everyday cognition. Yet, they provide a further level of abstraction beyond the object level and could be used in a straightforward fashion to participate in the downstream representations of supramodal concepts (see Discussion).

### Comparison between feature sets

To test our hypothesis, we compared the performance of each feature set for several small numbers of training examples (Fig 2). The results confirm the predictions: For small numbers of training examples, feature sets extracted later in the visual hierarchy generally outperformed features sets extracted earlier in the visual hierarchy. Critically, as predicted, we see that the conceptual features dramatically outperform generic_1_ features for small numbers of training examples (for 2, 4, 8, 16, and 32 positive examples). In addition, conceptual and generic_1_ features outperform generic_2_, which outperforms generic_3_. These results suggest that combinations of generic_1_ features are frequently consistent across small sets of examples without generalizing well to the entire category; patterns among categorical features, by contrast, tend to generalize much better for small numbers of examples.

**Fig 2.**
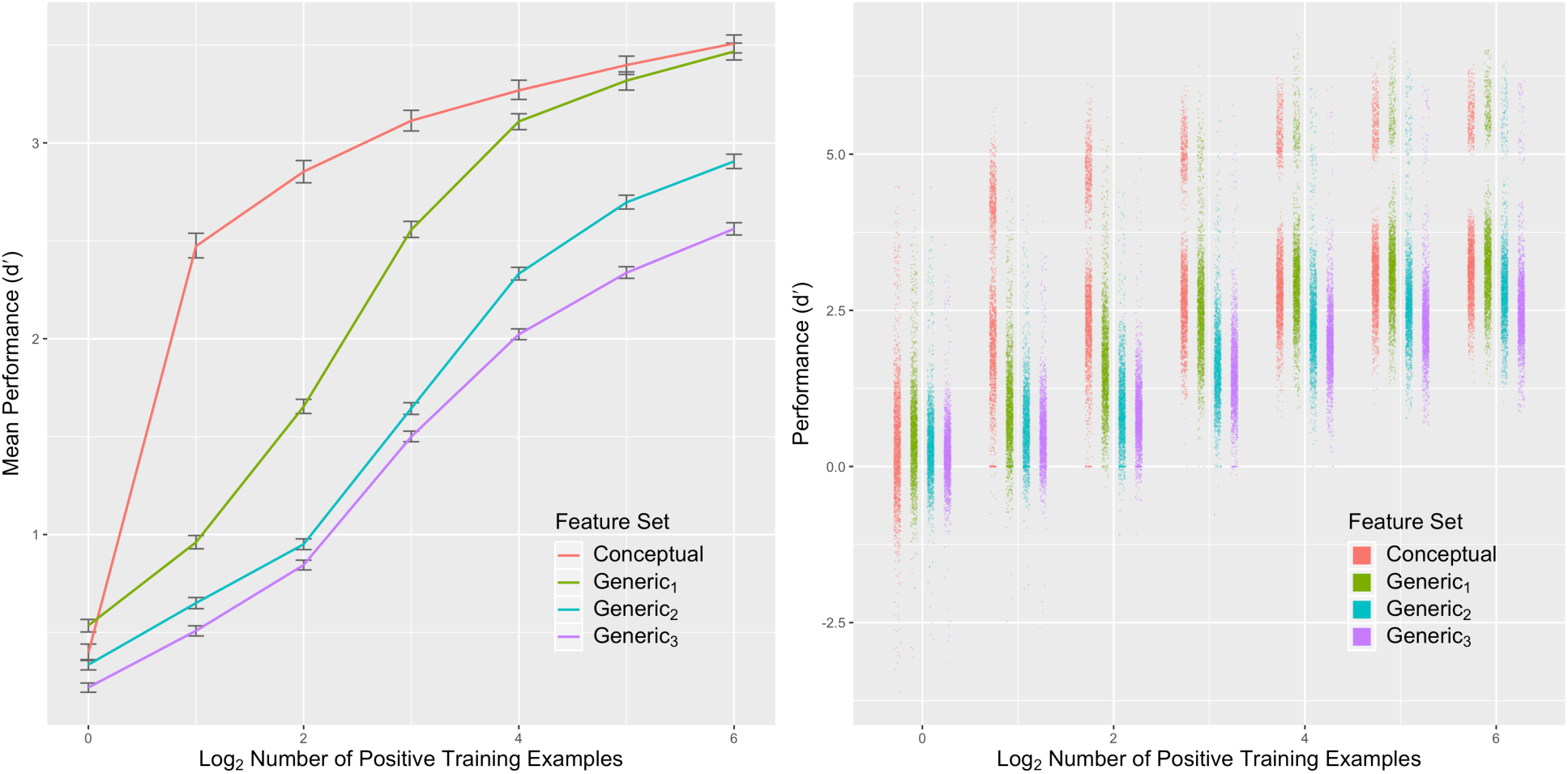
Main simulation performance summary. **Left**. Mean performance (y-axis) of classifiers in our analysis by feature set (color) and number of positive training examples (x-axis). **Right**. Performance (y-axis) of each classifier in our analysis (individual points) by feature set (color) and number of positive training examples (x-axis). Performance in both plots is measured as d′. Error bars are bootstrapped 95% CIs.

To verify this pattern quantitatively, we constructed a linear mixed effects model predicting d′ from main effects of number of training examples, and feature set, as well as an interaction between feature set and number of training examples, with a random effect of category. A Type III ANOVA analysis using Satterthwaite’s method finds main effects of feature set (*F*(3, 55,873) = 9105.5, *p* < 0.001) and number of training examples (*F*(6, 55,873) = 15,833.5, *p* < 0.001), as well as an interaction between feature set and number of training examples (*F*(18, 55,873) = 465.1, *p* < 0.001). We further find via single term deletion that the random effect of category explains significant variance (*χ*^2^(1) = 20,646.5, *p* < 0.001).

Having established a main effect of feature set, we further analyzed differences in performance between feature sets by computing pairwise differences in estimated marginal mean performance. Critically, we found that the conceptual features outperformed generic_1_, generic_2_, and generic_3_ features, generic_1_ outperformed generic_2_ and generic_3_ features, and generic_2_ outperformed generic_3_ (*p*s < 0.001).

The interaction between feature set and number of training examples is also supported by pairwise differences in estimated marginal mean d′. Critically, we find that conceptual features outperform the generic_1_ features for 2–32 positive training examples (*p*s < 0.001) and marginally outperform them for 64 positive training examples (performance difference = 0.041, p = 0.074). Thus, as predicted, leveraging prior concept learning leads to dramatic improvements in the ability of deep learning systems to learn novel concepts from few examples.

## Discussion and conclusion

A striking feature of the human visual system is its ability to learn novel concepts from few examples, in sharp contrast to current computational models of visual processing in cortex that all require larger numbers of training examples [30,31,44]. Conversely, previous models of visual category learning from computer science that perform well for small numbers of examples [47,48] (albeit not at the level of current state-of-the-art approaches) were not explicitly motivated by how the brain might solve this problem and do not provide biologically plausible mechanisms. It has been unclear, therefore, how the brain could learn novel visual concepts from few examples.

In this report, we have shown how leveraging prior concept learning can dramatically improve performance for few training examples. Crucially, this performance was obtained in a model architecture that directly builds on and extends our current understanding of how the visual cortex, in particular inferotemporal cortex, represents objects [30]: By using a “conceptual” layer, akin to concept representations identified downstream from IT in anterior temporal cortex [15,21,49,50] new concepts can be learned based on just two examples. This suggests that the human brain could likewise achieve its superior ability to learn by leveraging prior learning, specifically concept representations in ATL. How could this hypothesis be tested? In case disjoint neuronal populations coding for related concepts learned at different times can be identified, causality measures such as Granger causality [51–53] could provide evidence for their directed connectivity. At a coarser level, longer latencies of neuronal signals coding for more recently learned concepts relative to previously learned concepts would likewise be compatible with novel concept learning leveraging previously learned concepts.

Our scheme separates object-level and concept-level representations from task-specific decision mechanisms. As a result, object-based and conceptual representations can remain broadly tuned, supporting rapid learning for novel tasks. In fact, new concept learning was poor when using post-decision (i.e., post-softmax) units as input, whose responses were sharpened to exclusively respond to just a single concept. Our simulations therefore predict a key role for broadly tuned concept units, e.g., in the ATL, in enabling the rapid learning of novel concepts. Testing this hypothesis will require high-resolution methods like electrocorticography (ECoG, see, e,g., [54]).

Intuitively, the requirement for two examples to successfully learn novel concepts makes sense as this allows the identification of commonalities among items belonging to the target class relative to non-members. However, the phenomenon of Fast Mapping suggests that under certain conditions, humans can learn concepts even from a single positive and negative example. In contrast, in our system, performance for this scenario was generally poor. Yet, theoretically, one positive and one negative example should already be sufficient *if the negative example is chosen from a related category that would serve to establish a crucial, category-defining difference*, which is precisely what is done in conventional Fast Mapping paradigms in the literature. In the simulations presented in this paper, our negative example was chosen randomly, so we would not necessarily expect good ability to generalize from a single positive example. Yet, studying how variations in the choice of negative examples can further improve the ability to learn novel concepts from few examples would be a highly interesting question for future work that can easily be studied within the existing framework. In that context, an interesting observation in Fig 2b is that for the case of one positive and one negative example, the distribution for the conceptual feature set actually appears *bimodal*, with a number of very high d′ values (∼3) and many very low ones (∼0).

Another interesting question is whether there are conditions under which leveraging prior learning leads to suboptimal results compared to learning with features at lower level of the hierarchy, e.g., generic_1_. In particular, generic_1_ features are as good as conceptual features for larger numbers of training examples. Future work could explore whether there is some point at which features similar to generic_1_ outperform learning based on conceptual features: For instance, when lots of examples are available, does it help to learn the category boundaries directly based on shape rather than by relating the new category to previously learned ones?

Finally, while our work in this paper focused on visual concepts, it will be interesting to explore whether similar mechanisms are at work for even higher-level generalization, as in the learning of schemas. Work with rats shows that they can learn new paired associations between odor and place after just a single training trial, and an investigation of gene expression during this task suggests that learning new information leverages previously learned neocortical schemas [55,56]. These findings are consistent with fMRI studies showing heightened activity in medial prefrontal cortex for subjects learning information in a field they have already begun to study and heightened medial temporal activity for learning in an entirely novel field [57]. They are also consistent with neural network simulations of the Complementary Learning Systems theory of semantic memory that show fast learning for information consistent with an existing semantic network [58].

## Materials and methods

### Image Net

ImageNet (www.image-net.org) organizes more than 14 million images into 21,841 categories following the WordNet hierarchy [46]. Crucially, these images come from multiple sources and vary widely on dimensions such as pose, position, occlusion, clutter, lighting, image size, and aspect ratio. This image set has been designed and used to test large-scale computer vision systems [59], including models of primate and human visual object recognition [30,31]. We similarly use disjoint subsets of ImageNet to both train and validate a modified GoogLeNet and to train and test a series of binary classifiers.

To train and validate GoogLeNet, we randomly selected 2,000 categories from 3,177 ImageNet categories providing both bounding boxes and more than 732 total images (the minimum number of images per category in the Image Net Large Scale Visual Recognition Challenge (ILSVRC) 2015), thus ensuring each category represented a concrete noun with significant variation (S1 Table). One of the authors further reviewed each category to ensure it represented a concrete visual category. We set aside 25 images from each category to serve as validation images and used the remainder as training images. We thus used a total of 2,401,763 images across 2,000 categories for training and 50,000 images across those same 2,000 categories for validation. To reduce computational complexity, all images were resized to 256 pixels on the shortest edge while preserving orientation and aspect ratio and then automatically cropped to 256 x 256 pixels during training and validation. While it is possible for this strategy to crop the object of interest out of the image, previous work with the GoogLeNet architecture [45] suggests that the impact on performance is marginal.

To train and test our binary classifiers, we used the training and validation images from 100 of the 1,000 categories from the ILSVRC2015 challenge [59]. As with the GoogLeNet images, all images were resized to 256 pixels on the shortest edge while preserving orientation and aspect ratio and then automatically cropped to 256 x 256 pixels during feature extraction.

### GoogLeNet

GoogLeNet is a high-performing [45] deep neural network (DNN) designed for large-scale visual object recognition [59]. Because prior work has shown that the performance of DNNs is correlated with their ability to predict neural activations [29,30] and that GoogLeNet in particular is a comparatively good predictor of neural activity [31], we use GoogLeNet as a model of human visual object recognition. Because the exact motivation for GoogLeNet and the details of its construction have been reported elsewhere, we focus here on the details relevant to our investigation. We used the Caffe BVLC GoogLeNet implementation with one notable alteration: we increased the size of the final layer from 1,000 to 2,000 units, commensurate with the 2,000 categories we used to train the network. We trained the network for ∼133 epochs (1E7 iterations of 32 images) using a training schedule similar to that in [45] (fixed learning rate starting at 0.01 and decreasing by 4% every 3.2E5 images with 0.9 momentum), achieving 44.9% top-1 performance and 73.0% top-5 performance across all 2,000 categories.

### Main simulation

To study how previously learned visual concepts could facilitate the learning of novel visual concepts, we trained a series of one-vs-all binary classifiers (elastic net logistic regression) to recognize 100 new categories from the ILSVRC2015 challenge. The 100 categories were chosen uniformly at random and remained constant across all feature sets (S2 Table).

The primary hypothesis of this paper is that prior learning about visual concepts can significantly improve learning about new visual concepts from few examples. Learning new categories in terms of existing category-selective features is thus of primary interest, so we compared several feature sets to test the effectiveness of learning from category-selective features relative to other feature types. We specifically compared the following feature sets:

- Conceptual: 2,000 features extracted from the `loss3/classifier`, a fully connected layer of GoogLeNet just prior to the softmax operation producing the final output.
- Generic_1_: 4,096 features extracted from `pool5/7×7_s1`, an average pooling layer of GoogLeNet (kernel: 7, stride: 1) used in computing the final output.
- Generic_2_: 13,200 features extracted from the `loss2/ave_pool`, an average pooling layer of GoogLeNet (kernel: 5, stride: 3) mid-way through the architecture used in computing a second training loss.
- Generic_3_: 12,800 features extracted from the `loss1/ave_pool`, an average pooling layer of GoogLeNet (kernel: 5, stride: 3) early the architecture used in computing a third training loss.
- Generic_1_ + Conceptual: 4,096 Generic_1_ features combined with 2,000 Conceptual features for a total of 6,096 features.

All features were selected for broad tuning to encourage generalization. The Conceptual features — being as close to the final output as possible but without the task-specific response sharpening of the softmax operation — represent what should be the most category-sensitive features of GoogLeNet (i.e., individual features serve as more reliable signals of category membership than features from other feature sets; S3 Appendix). The various Generic feature sets were chosen as controls against which to compare the conceptual features. Based on prior work using GoogLeNet, these layers likely correspond to high-level visual cortex (e.g. V4, IT, fusiform cortex) [30,31]. The Generic_1_ features act as close controls against which to compare the conceptual features. These features provide a representative basis in which many visual categories can be accurately described while themselves being relatively category-agnostic (S3 Appendix). We chose a layer near the end of the network but before the fully connected layers that recombine the intermediate features into category-specific features. The GoogLeNet architecture defines two auxiliary classifiers—smaller convolutional networks connected to intermediate layers to provide additional gradient signal and regularization during training—at multiple depths in the network. We define the Generic_2_ and Generic_3_ features using layers from these auxiliary networks that correspond to the layer from the primary classifier used to define Generic_1_.

We measured feature set performance by training a series of one-vs-all binary classifiers (elastic net logistic regression) for each feature set, meaning that each feature set served in a sub-simulation as the sole input to the classifiers. For each feature set, we trained 14,000 classifiers— one for each combination of test category, number of training examples, and random training split—and measured performance using d′. Our ImageNet ILSVRC-based image set had 100 categories (See ‘ImageNet’ above). Positive examples were randomly drawn from the target category, while negative examples were randomly drawn from the other 99 categories. Because we were interested in how prior knowledge helps with learning from few examples, we tested classifiers trained with *n* ∈ {2, 4, 8, 16, 32, 64, 128} total training examples, evenly split between positive and negative examples. To better estimate performance and average out the effects of the classifiers’ random choices, we repeated each simulation by generating 20 random training/testing splits unique to each combination of test category and number of training examples.

## Acknowledgements

The authors thank Jacob G. Martin for helpful conversations and Benjamin Maltbie for help with running simulations.

## Supporting information

**S1 Table. 2**,**000 ImageNet categories used to train the GoogLeNet object recognition network.** A comma-delimited table listing the WordNet ID, a short natural-language title, and a short natural language gloss for each of the 2,000 categories used to train the modified GoogLeNet object recognition network used in this paper.

**S2 Table. 100 ImageNet categories used to compare feature sets.** A comma-delimited table listing the WordNet ID, a short natural-language title, and a short natural language gloss for each of the 100 categories used to compare feature sets extracted from GoogLeNet.

**S3 Appendix. Category selectivity analysis.** An additional analysis showing that, for the four feature sets examined in this paper, the closer a feature set is to the final output of the network, the more category-selective that feature set is (i.e., individual features more reliably signal category membership).

