## Supplementary material for "Leveraging prior concept learning improves ability to generalize from few examples in computational models of human object recognition": S3 Appendix

### S3 Appendix. Category Selectivity Analysis

#### Results

We hypothesized that the closer in the processing hierarchy a feature set was to the final output of the network, the more category-selective its features would become. By category-selective, we mean that individual features provide a reliable signal of category membership (*i.e.*, individual features can accurately classify images as positive or negative examples of the category). To test this prediction, we compared the classification performance of individual features from the four feature sets for the 100 categories used in the main simulation (Fig. A).

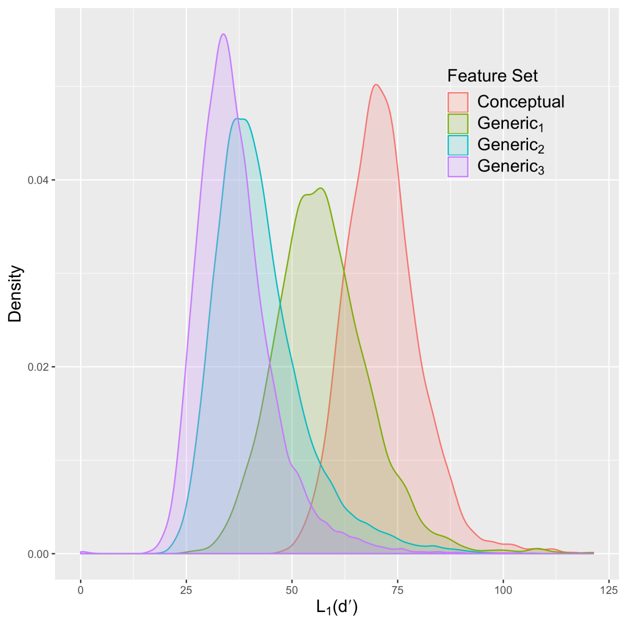


**Figure A. Category Selectivity.** Gaussian kernel density estimate (y-axis) of single feature d′ (x-axis) by feature set (individual curves). d' of each feature in each feature set is the $L_1$ norm computed over 100 test categories.

We were interested in assessing the full category selectivity of each feature across a variety of categories. In the following, we therefore specifically analyze the $L_1$ norm over d′ classification scores for each feature across the 100 categories used in the main simulation. This approach still provides thousands of data points per feature set.

Qualitatively, we find the predicted pattern: individual features extracted later in the visual hierarchy outperformed (in the sense of having higher $L_1$ norms and thus higher overall selectivity) feature sets extracted earlier in the visual hierarchy. We specifically see that the conceptual features outperform $\text{generic}_1$, which outperforms $\text{generic}_2$, which outperforms $\text{generic}_3$.

To verify this pattern quantitatively, we constructed a linear mixed effects model predicting d′ from a main effect of feature set, with a random effect of feature. A Type III ANOVA analysis using Satterthwaite's method finds main effects of feature set ($F(3, 22998.6) = 10196.6$, $p < 0.001$), and single term deletion finds that the random effect of feature also explains significant variance (category: $\chi^2(1) = 156.4$, $p < 0.001$).

Having established a main effect of feature set, we further analyzed differences in performance between feature sets by computing pairwise differences in estimated marginal mean performance. Critically, we found that: conceptual features outperformed $\text{generic}_1$, $\text{generic}_2$, and $\text{generic}_3$ features; $\text{generic}_1$ outperformed $\text{generic}_2$ and $\text{generic}_3$ features; and $\text{generic}_2$ outperformed $\text{generic}_3$ ($p$s $< 0.001$).

**Method**

To better understand the differences between the feature sets tested in the main simulation, we trained a second series of one-vs-all binary classifiers to recognize the 100 categories used in the main simulation. The main simulation assumed that the features of the conceptual feature set were more category-selective than the features of the other available feature sets, and more generally, that features extracted later in the visual hierarchy would be more category-selective than features extracted earlier in the visual hierarchy. By category selective, we mean that individual features provide a reliable signal of category membership (i.e., individual features can accurately classify images as positive or negative examples of the category). To test this assumption, we compared individual features from each of the four feature sets in the main simulation, namely conceptual, $generic_1$, $generic_2$, and $generic_3$.

Because our interest in this analysis is specifically on measuring the classification performance of individual features, we trained a one-vs-all binary classifier (elastic net logistic regression) for each combination of feature and category. Each individual feature from the four tested feature sets thus served as the sole input to one of the trained classifiers. Our ImageNet ILSVRC-based image set had 100 categories (See ‘ImageNet’ in main text). Positive examples were randomly drawn from the target category, while negative examples were randomly drawn from the other 99 categories. Our interest in this analysis was less in learning from few examples than in understanding a feature’s category-selectivity; for each classifier, we thus randomly selected 64 positive and 64 negative examples as training examples. Our test set contained an unbalanced number of positive and negative examples, so we measured performance using $d'$.
